# mCytoCounter, a deployable somatic cell counting device for the assessment of mastitis in dairy farms

**DOI:** 10.1101/2025.11.19.689399

**Authors:** Neha Khaware, Tushar Jeet, Prabhu Balasubramanian, Holger Schulze, Vivekanandan Perumal, Till T Bachmann, Ravikrishnan Elangovan

**Author notes:** Corresponding author: Ravikrishnan Elangovan, Professor, Department of Biochemical Engineering & Biotechnology, Indian Institute of Technology Delhi, New Delhi, India.

## Abstract

Bovine mastitis is a recurring disease among herds that is often caused by unhygienic farm conditions, resulting in bacterial invasion of the udder. It is a leading cause of financial losses for dairy farmers, largely due to the widespread use of antimicrobials for treatment, which in turn fuels the development of drug-resistant bacteria. Somatic cell count (SCC) is a widely used quantitative method for diagnosing sub-clinical and clinical mastitis. We have developed a rapid and quantitative microfluidic device that enables the quantification of somatic cells in milk, thereby aiding in the diagnosis of bovine mastitis. Our platform features a microfluidic cartridge for capturing somatic cells, which is inserted into an in-house device called the mCytoCounter. The device reads the fluorescence emission signals from the somatic cells captured on the membrane in 2 minutes, with a total turnaround time of 15 minutes. The mCytoCounter is an automated and deployable system that can be used in dairy farms with minimal training. We analysed the performance of mCytoCounter with milk samples and benchmarked it against the SCC using a fluorescence microscope with a correlation coefficient of 0.97. The mCytoCounter has a detection range of 10^3^ to 10^6^ SCC/mL, which covers the required cell number range to discriminate between healthy, subclinical, and clinical mastitis.

## 1. Introduction

Bovine Mastitis is the most common disease among lactating dairy animals, resulting in low production and inferior-quality milk. Mastitis is mainly caused by a bacterial invasion, resulting in the influx of leukocytes, accompanied by inflammation of the udder, and results in an increase in the SCC in the produced milk. Hence, SCC in milk is an indicator of bovine health and is an early biomarker for subclinical and clinical mastitis. Globally, there are an estimated 270 million dairy cows (WWF, 2022). India is the world’s largest milk producer, with a population of 57 million dairy cows, which produced 239.30 million metric tonnes of milk in the fiscal year 2023-2024 (1). The economic loss due to mastitis every year is tremendously high. Uttar Pradesh alone in India suffers a yearly loss of 157.51 million USD due to subclinical and clinical mastitis (2). In India, the economic losses from mastitis have increased by about 115-fold in the last five decades and cost about 860 million USD annually (3).

Mastitis is majorly a result of intramammary infection, defined as the presence of a pathogen in the mammary gland resulting in milk contamination, making it unfit for consumption(4). Somatic cells are a mixture of milk-secreting epithelial cells and white blood cells (WBCs) (80%) that enter the mammary gland(5). Typically, <200,000 SCC/mL is considered uninfected, 300,000 to 500,000 SCC/mL is subclinical mastitis, and >500,000 SCC/mL (6). In clinical mastitis cases, the SCC can go up to >1X10^6^ cells/mL (7). Clinical mastitis is inferred from the presence of visibly abnormal milk and inflamed udder and is mostly evaluated by SCC, fever and inflammation of the udder (5). Subclinical mastitis is interpreted by the presence of inflammation but visibly normal milk and mammary glands. Due to no visible depletion in the milk quality, it often goes unnoticed, as farmers can’t detect it without tests like SCC. This makes it a bigger problem as it can silently spread in the herds before being detected, resulting in a cumulative effect on the yield and herd health (8). Some common tests evaluated for diagnosis of subclinical mastitis are SCC, lactose percentage, measurement of lactate dehydrogenase activity, and electrical conductivity (5,9,10).

Majority of bovine mastitis is caused by *Staphylococcus aureus* and it spreads easily in the herd if left untreated (Patel et al., 2021). The reported prevalence of *S. aureus-induced* mastitis, however, varies considerably depending on the form of the disease and geographical location. A comprehensive review noted that *S. aureus* can account for approximately 10.6% of clinical cases and a much higher 29.1% of subclinical mastitis (12). A regional study from India reported an even higher prevalence of 54% in clinical and 50% in subclinical mastitis (13). Other than *S. aureus,* different species of *Staphylococcus* (45%)*, E. coli* (14%), *Streptococcus* spp. (13%), *Mycobacterium bovis, Klebsiella* are responsible for causing sub-clinical and clinical mastitis in India (14) (15). High numbers of antibiotic resistance rates of these bovine mastitis-causing pathogens, including many multidrug-resistant isolates, have been reported in various countries in recent years (16–19). These antimicrobial-resistant strains can be transferred to other cows, buffaloes in the herd and humans through milk and meat. High levels of *S. aureus* resistance to most antibiotics, such as penicillin, erythromycin, tetracycline, gentamicin, tobramycin, kanamycin, and methicillin (MRSA), making it especially difficult to contain. Due to economic pressure, antibiotic consumption is high, leading to the selection of AMR strains (20). The early detection of mastitis can reduce the incidence of severe cases of intramammary infection and, consequently, reduce antimicrobial usage. The need for a rapid and economical test that can help dairy farmers diagnose bovine mastitis at an early stage is increasingly recognized.

Some widely used methods for estimating somatic cells in milk are-direct SCC, California mastitis test (CMT), Wisconsin mastitis test (WMT), and esterase activity test. There are numerous automated somatic cell counters that utilise microscopy for cell counting (21). Most of them use fluorescent dye or a stain to label the cells and visualize them under a microscope, followed by manual or automated counting. There are reference methods for cell counting, such as flow cytometry (22). However, these tests are only available at dairy veterinary clinics, reducing their utility as an on-farm diagnostic tool. Also, these counting methods are either expensive, time-consuming, or require manual intervention. Quantitative fluorescence assays can help eliminate the manual counting steps, saving time per reaction. CMT, WMT, and esterase activity tests are either qualitative or semi-quantitative on-farm (cow-side) tests used to estimate SCC in milk (23).

Microfluidics has emerged as a powerful tool in various applications, ranging from basic biological research to clinical research and development (24). Microfluidics is a system that enables the processing and manipulation of small volumes of liquids in channels with dimensions in micrometres. It has many advantages, such as the ability to use a small amount of sample, improved resolution, and sensitivity in separation and detection, reasonably low cost compared to available reference methods, and shorter analysis time (25). Separation of a specific cell type from a complex sample containing a heterogeneous cell population has many applications, especially in biomedical and biochemical research (26).

A vast array of active and passive methods has been employed to separate blood cells. Active methods constitute technologies that operate based on an external field or a force, while passive methods use the physicochemical properties of the cells (27). Micropillar structures, microporous membranes, etc., can separate cells and beads based on their size and compressibility, which can be exploited to separate WBCs from erythrocytes or plasma from the rest of the cells. Although many have reported that these types of separations may result in clogging, that can be overcome using proper dilutions (Miao Ji et al., 2008),(29).

Conventional cell sorting techniques include flow cytometry, Fluorescence-activated cell sorting (FACS), and Magnetic-activated cell sorting (MACS). These methods offer excellent specificity and high-throughput screening with substantial data outputs. However, they are not widely used in clinical settings due to the following factors: higher instrument cost, intensive training of the user personnel, higher initial and operating costs in labeling antibodies, sheath fluid, magnetic nanoparticle, and magnetic capture-based systems (30),(31).

Here, we present a microfluidic cartridge and a reader capable of quantifying somatic cells in the milk with a short turnaround time, in farm settings. The somatic cells from milk are enriched on the membrane and washed to remove any background signal. We have benchmarked the performance of mCytocounter against conventional imaging-based SCC and shown its utility for farm deployment.

## 2. Materials and Methods

The entire assay is performed in a 3D-printed microfluidic cartridge and device with a temperature controller, fluorescence reader and control electronics. The device can evaluate two cartridges simultaneously. Each cartridge has four channels and can be used to read four different milk samples.

### 2.1 Cartridge and Device Fabrication

The cartridge was 3D printed with a ProJet MJP 2500 [3D Systems] using VisiJet MJP 2500-CL and WH (white, translucent material). It has dimensions of 70 x 40 x 5mm (L x B x H). The cartridge has four sample wells that can accommodate 50µL of the sample. The cartridge assembly has four components-top sheet, membranes, absorbent pad for collecting dump liquid and bottom sheet to vacuum pack the cartridge as explained in the supplementary figure_S4. The sample can be added to the cartridge well using a pipette or a syringe. The sample then flows through the serpentine channel and crosses the membrane. The somatic cells are captured on the glass fibre B membrane [GE Healthcare Life Sciences, Whatman^TM^, 1821-047], and the pass-through liquid is collected in the dump chamber. The device is an in-house, fully automatic fluorescence reader equipped with a vacuum pump to control liquid flow in the microfluidic cartridge. The device has a touchscreen pad that will allow users to control the system with the help of a Graphical User Interface (GUI) (figure 1A, 1B).

**Figure 1.**
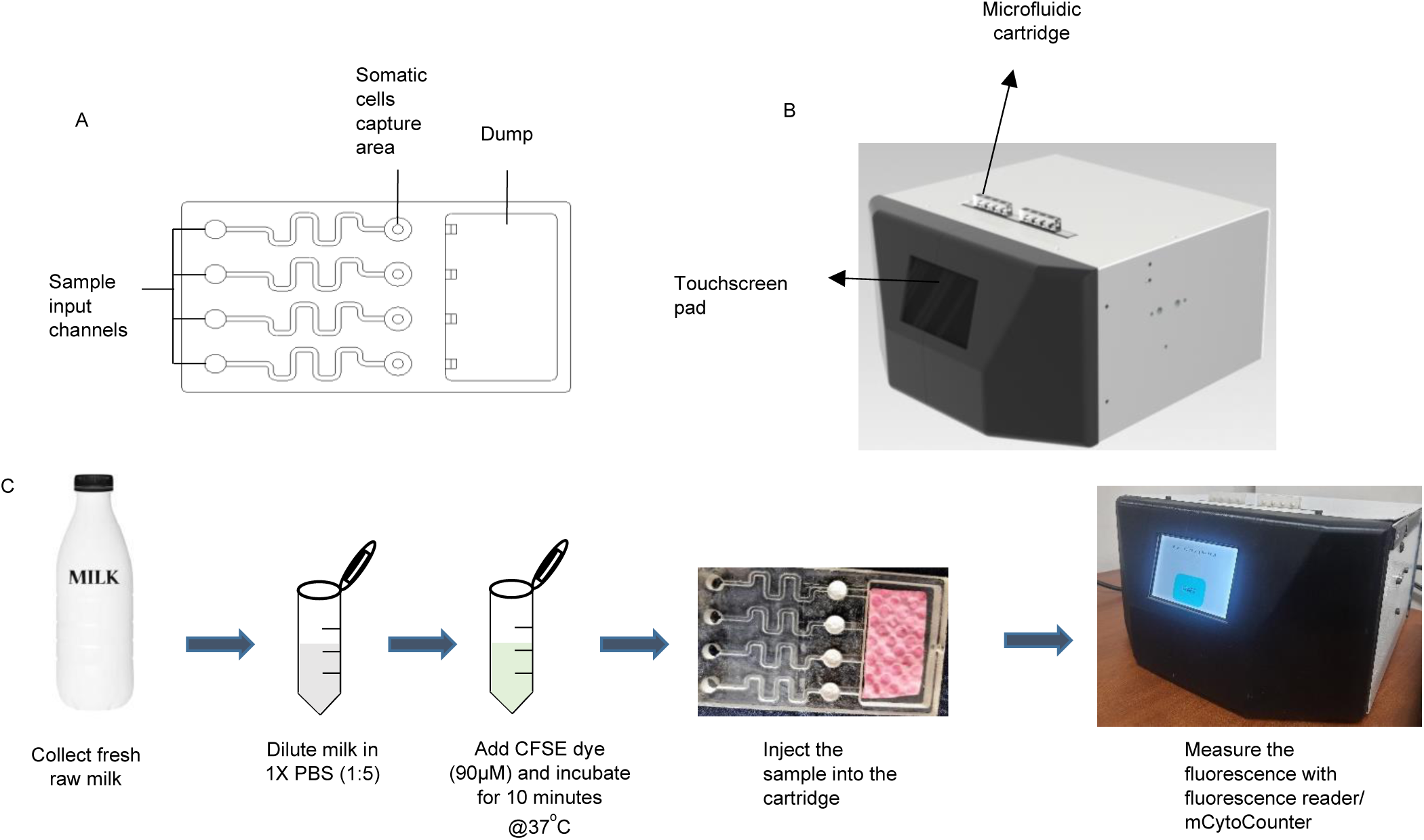
A. Cartridge, B. mCytoCounter device, C. Assay protocol

### 2.2 Fluorescent reader

The optical system is a customized setup consisting of an LED, lenses, excitation and emission filters, and a dichroic mirror. A spherical lens with a focal length of 12mm is combined with an LED and a filter with an excitation wavelength of 480 nm and an emission wavelength of 510 nm (long pass). The detector is a photodiode integrated with in-house software to calculate fluorescence signal in the form of Relative Fluorescence Units (RFU). The values obtained can be compared with the calibration graph to get the approximate cell count. The fluorescence reader is mounted on a linear guide allowing independent reading of fluorescent signals from two different cartridges and eight wells. The process flow of the mCytoCounter is shown in Figure 2B. The electronic assembly ensures a fully automatic data acquisition and GUI. Supplementary figure_S2 shows the block diagram of the printed circuit board. The hardware has been designed in a modular fashion where the microcontroller, an integrated circuit interacts with the GUI via touchscreen pad and initiates the program.

**Figure 2.**
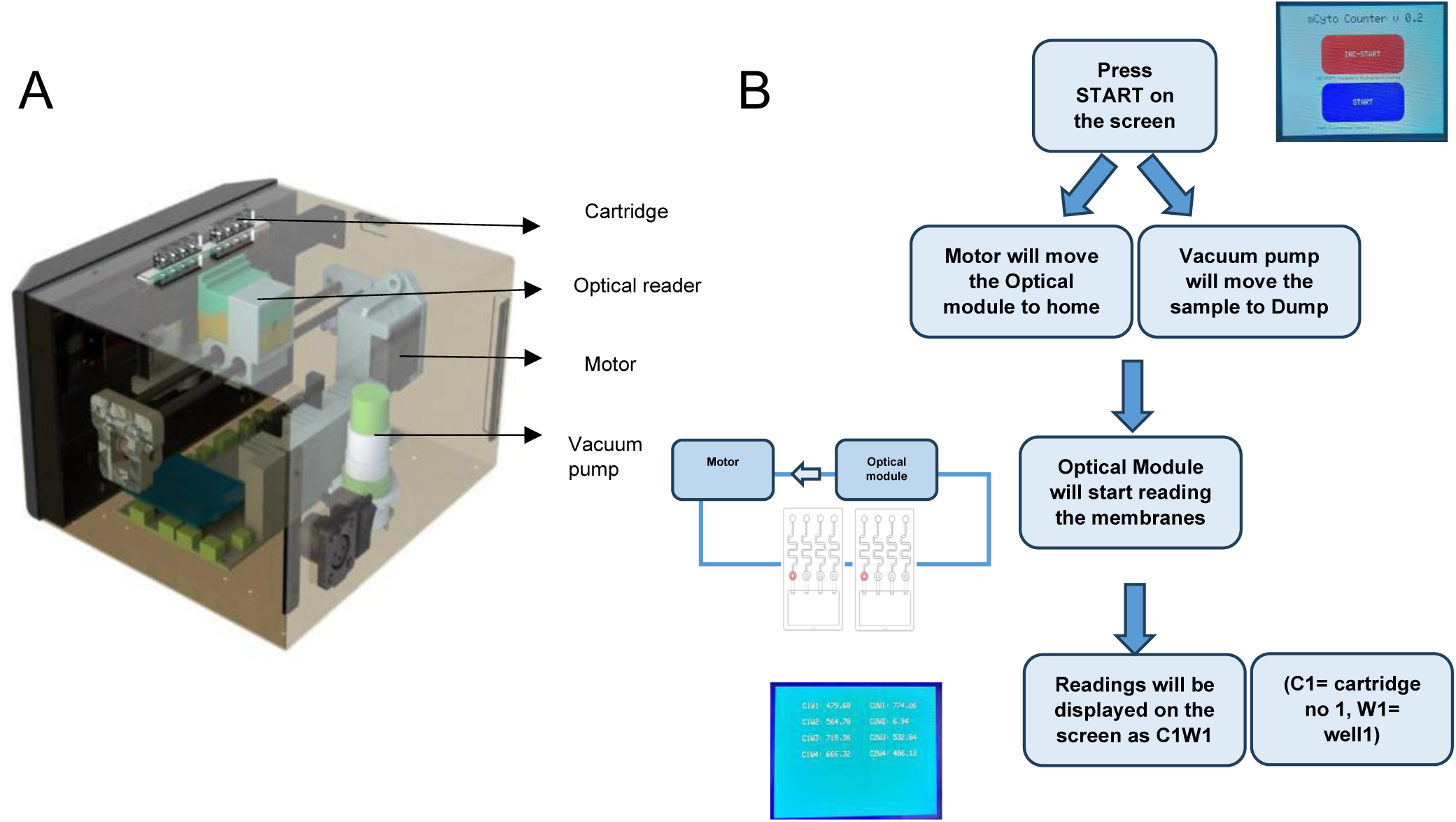
Components of mCytoCounter. A. 3D view of mCytoCounter, B. Flow diagram of mCytoCounter working. C1= Cartridge no 1 and W1= Well 1

### 2.3 Experimental approach

#### 2.3.1 Spiked samples

The somatic cells were isolated from whole cow milk using centrifugation (Supplementary Figure_S1). Briefly, the milk (40 mL) was collected in a falcon tube and centrifuged at 1800g for 30 minutes. The supernatant was discarded, and the pellet was washed two times with 1X PBS [Sigma, P4417-100TAB] at 1500g for 10 minutes. The final pellet was then resuspended in 1mL PBS. The isolated somatic cells were counted using a hemocytometer [Rohem, India]. For ease of counting, the cells were labeled with methylene blue in a ratio of 0.1:4 (Methylene blue: Cells). The known number of somatic cells was then used to prepare the calibration curve with different dilutions. The dilutions were prepared in depleted milk (Milk without cells; removed by centrifugation) to mimic the actual conditions. The plot was made using *OriginPro* 2020b, data analysis, and graphing software.

#### 2.3.2 Direct Milk Samples

Whole milk samples were diluted five times with 1X Phosphate Buffered Saline (PBS) (1:5) [Sigma, P4417-100TAB] and labeled with the CFDA, SE (5-(and-6)-Carboxyfluorescein Diacetate, Succinimidyl Ester) also called CFSE dye [ThermoFisher, C1157] for 10 minutes at 37°C to fluorescently label the somatic cells. About 80µL of the sample is added into the cartridge with the help of a pipette into the sample input channel. The sample is moved to the filter by a vacuum pump installed in the m-CytoCounter. The filter captures the somatic cells and lets the remaining liquid pass to the dump. The filter is a glass microfiber-B (GF/B) with a pore size of 1µm, laser-cut to a diameter of 5mm to fit into the cartridge. The captured cells on the filter are then excited, and the emission signal is read by an in-house fluorescent reader installed in the device. The entire assay, from sample preparation and addition into the cartridge to the fluorescence readout, takes about 15 minutes (Figure 1C). The plot was created using OriginPro 2020b, a data analysis and graphing software.

## 3. Results and discussion

### 3.1 Evaluation of Membranes and dyes

The somatic cell capture and labeling were evaluated with five different filter membranes and four labeling dyes, respectively. This was done to optimize the cell retention in the cartridge with uniform labeling of the cells thereby ensuring better reproducibility of the assay.

Five different filter membranes were evaluated for the performance with somatic cell capture (Figure 3A). Glass Fibre-B, F, D, CP and polycarbonate track-etched membrane (PCTE) were used [GF/F, GF/D, GF/CP, and PCTE were purchased from GE Healthcare Life Sciences Whatman^TM^ 1825-047, GE Healthcare Life Sciences Whatman^TM^ 1823-047, Sigma GFCP001000, Cytiva Whatman^TM^ 10418306, respectively]. The blank membranes and membranes with captured cells were evaluated under an inverted fluorescence microscope (Olympus IX71). The somatic cells were labeled with CFSE dye (90 µM). The excitation and emission wavelengths for CFSE dye are 492nm/517nm. The Glass Fibre B (GF/B) performed the best out of all the membranes. It has a pore size of 1µm, which captures the maximum number of somatic cells without clogging due to its high porosity. Other glass fibre membranes had a higher background in blank membranes. Although the polycarbonate track-etched membrane also yielded comparable results, it has a low porosity (10%), leading to clogging and uneven cell capture (32).

**Figure 3.**
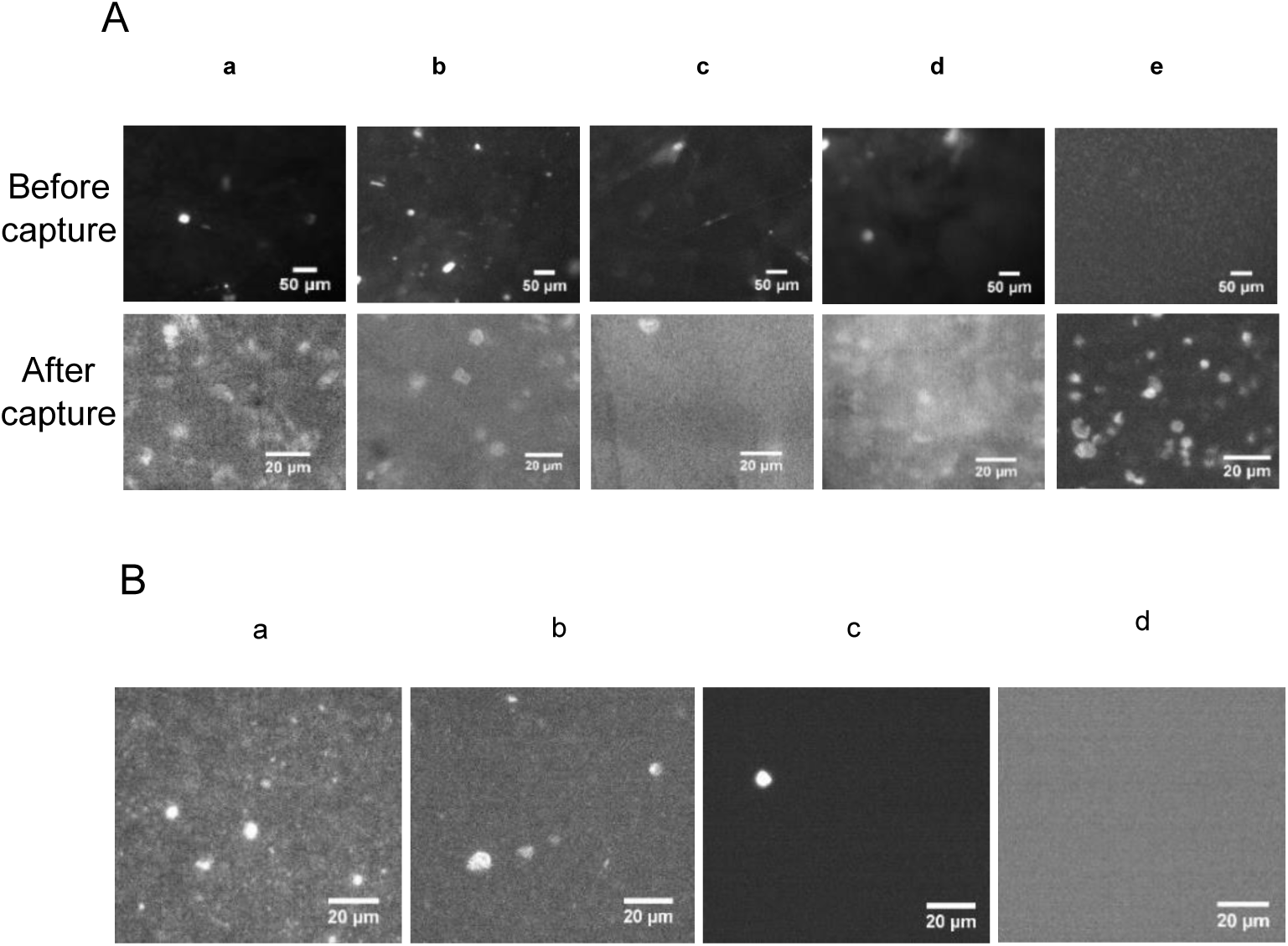
Evaluation of membranes and dyes. **A.** Different filters (used for capturing somatic cells) under a fluorescence microscope, blank (top) and with somatic cells (bottom). a= Glass Fibre B (GF/B), b= GF/D, c= GF/CP, and d= GF/F and e= Polycarbonate track-etched (PCTE). **B**. Evaluation of different labeling dyes. a= Acridine orange, b= CFSE, c= Eva Green and d= FDA. Sample is somatic cells spiked in milk. The cells were observed in a fluorescence microscope (Olympus IX71) cells to release a fluorescent product. It exhibited specific binding to somatic cells, with minimal or no binding to fats. Eva green dye is a cell-impermeable nucleic acid-binding dye and was used with Triton X-100 detergent. It was inefficient in binding to all the somatic cells, as the permeability due to the detergent was not uniform. Different concentrations of Triton X-100 were used (0.1%, 0.5%, 0.8%, and 1%) with EvaGreen, but none of the combinations could label somatic cells uniformly. FDA is a cell-permeable esterase-reacting dye, but it was incapable of binding to all the somatic cells. Therefore, CFSE dye was selected as the somatic cell labeling dye as it demonstrated superior performance compared to the other dyes evaluated.

The somatic cells were labeled using 4 different dyes-acridine orange, CFSE (5-(and-6)-Carboxyfluorescein Diacetate, Succinimidyl Ester), Eva green and FDA (fluorescein diacetate). The results obtained are shown in Figure 3B. Acridine orange was added to the somatic cell sample, spiked in milk and visualized under the microscope after 5 minutes. It is a cell-permeable nucleic acid binding dye, but resulted in higher background fluorescence signals as compared to other dyes. A lot of non-specific binding with fats was observed. CFSE is also a cell-permeable dye that reacts with esterases in viable

The GF/B membrane was also evaluated for capture performance using a multimode microplate reader (Victor Nivo, PerkinElmer). The first bar corresponds to the blank membrane, 2^nd^ bar is the membrane with captured 10^5^ cells/mL, 3^rd^ bar corresponds to the membrane with 10^6^ cells/mL (Figure 4). Also, no cells were leaked after the washing step as observed in the dump liquid.

**Figure 4.**
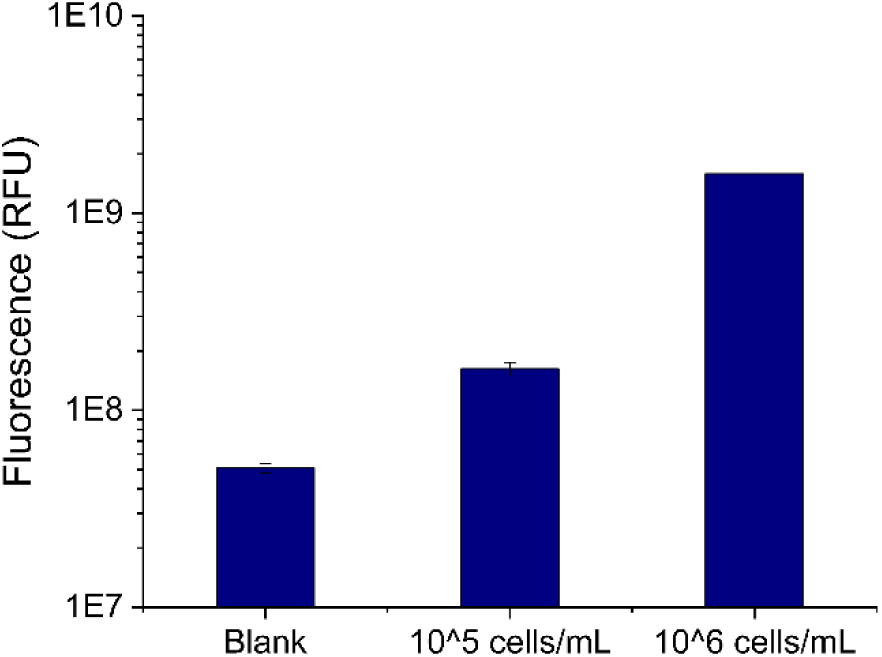
GF/B membrane evaluation; n = 3

### 3.2 Somatic cell counting in the microscope

The somatic cells were quantified in a fluorescence microscope (Supplementary Figure_S3). A calibration plot was developed using the known concentration of cells. The cell concentration was determined in a hemocytometer and was serially diluted in four dilutions ranging from 10^7^ to 10^4^ cells/mL. The dilutions were not made beyond 10^4^ cells/mL as it was difficult to spot cells in these diluted samples. The sample was added in a flow cell with dimensions 1cm X 1.8cm X 0.003cm (l*b*h). The average number of cells was calculated by counting the cells in 20 different frames using a 10x objective in a fluorescence microscope (Olympus IX71). The experiment was repeated thrice, and the average number of cells was then used to prepare a calibration plot (Figure 5A). The average cell count per field of view followed a linear fit over a four-log dynamic range. At lower concentrations, such as lower than 10000 cells/mL (∼1 to 2 cells/ frame), it becomes laborious to count cells and has a higher variation probability; hence, quantification from mCytoCounter would be helpful (supplementary Figure_S5).

**Figure 5.**
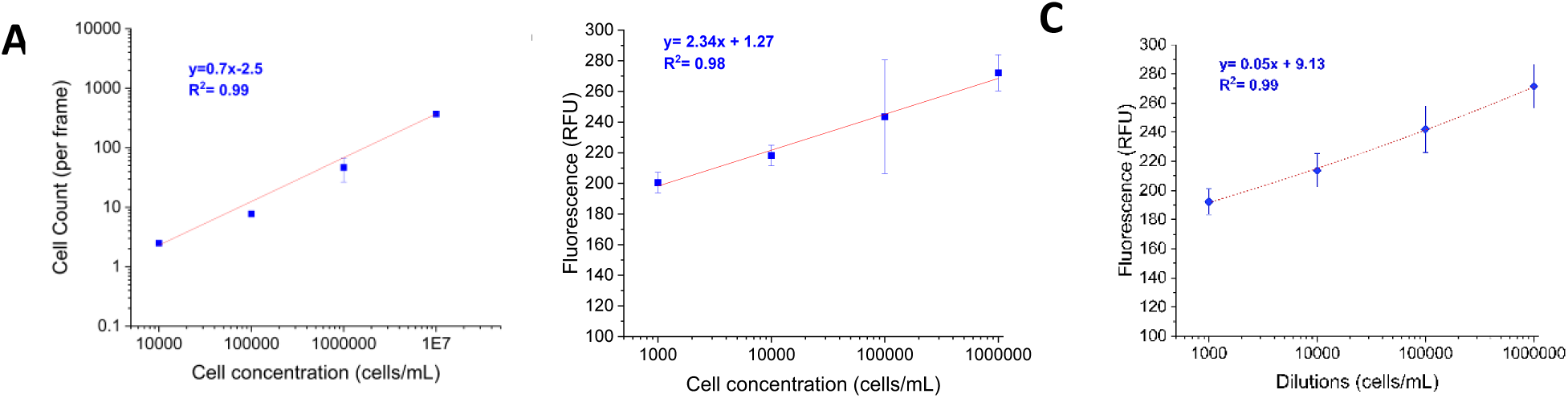
Calibration curves. **A**-Calibration curve from microscope. The experiment was repeated 4 times. **B**-Results in the column. The experiment was repeated 5 times. **C**-Results in the cartridge. The experiment was repeated 5 times. The somatic cells were isolated and spiked in the depleted milk (1:5). Error bars represent standard deviation

### 3.3 Somatic Cell Calibration (spiked samples)

Somatic cells isolated from milk have a heterogeneous population of cells consisting of epithelial cells and leukocytes(4,7,22). These cells differ in size, shape, and deformability. Epithelial cell diameter, for instance, ranges from 1-15µm(33,34).

The somatic cells were isolated from 80mL milk as explained in Supplementary Figure_S1. The isolated somatic cells were then counted using a hemocytometer and serially diluted in the diluted (1:5) depleted milk from 10^6^ to 10^3^ cells/mL. The calibration plot was produced using somatic cells spiked in diluted milk (1:5). The milk used for preparing dilutions was depleted (obtained after one round of centrifugation 1800g for 30 minutes) to remove fats and cells. The fluorescence was measured using a plate reader and an in-house mastitis diagnosis device (mCytoCounter) and recorded as Relative Fluorescence Units (RFU). The calibration plot for somatic cells spiked into the depleted milk has been displayed in Figure 5, where the serially diluted input sample was measured for fluorescence using the plate reader. For fluorescence measurement in the plate reader, the GF-B membrane in the cartridge were removed and checked directly in the multimode microplate reader (Victor Nivo, PerkinElmer) by placing it in a 96-well plate and obtaining readings.

### 3.4 Centrifugal enrichment versus mCytoCounter

The performance of mCytoCounter was evaluated using three different techniques. Firstly, the GF/B membrane was evaluated for its capture and fluorescence performance in a commercial plate reader for which the somatic cells were captured on the membrane in a column using centrifugation. Secondly, the cartridge was evaluated for its cell capture efficiency, and fluorescence was recorded in the commercial plate reader. Finally, the somatic cells were captured on the membrane in the cartridge and fluorescence was recorded in the mCytoCounter.

#### 3.4.1 Filtration using centrifuge and reading in the plate reader

The GF/B membrane was initially evaluated in the Qiagen QIAprep spin miniprep column (removed the Qiagen membrane and placed GF/B) for its ability to capture somatic cells. The membrane was carefully placed inside the column and fixed with O-rings. Somatic cells were diluted from 10^3^ to 10^6^ cells/mL in depleted milk. The labeled somatic cells sample was added from the top, and the columns were centrifuged at 300g for 10 minutes. The membrane was then removed from the column, and the emission signals were read using a multimode plate reader (PerkinElmer, Victor Nivo). The experiment was repeated five times and plotted as shown in Figure 5B. This experiment was conducted to verify that the GF/B membrane could efficiently capture somatic cells and that these cells could be detected for emission signals using a commercial plate reader.

#### 3.4.2 Filtration in the cartridge and reading in the plate reader

After evaluating the GF/B membrane in the column, the somatic cells were captured in the cartridge. The serially diluted somatic cells were externally labelled with CFSE and added to the cartridge. The liquid operations were performed manually using a syringe. As the sample moved in the cartridge, the somatic cells got captured in the cartridge and the remaining liquid moved to the dump. The membranes were removed from the cartridge and read in a multimode plate reader (PerkinElmer, Victor Nivo). This confirmed that the cartridge could capture the somatic cells, as no cells were observed in the dump liquid when observed under the microscope. The experiment was repeated five times and plotted as shown in Figure 5C. The coefficient of determination (R^2^) was 0.99, indicating excellent correlation and high reproducibility in lower cell concentrations, as well as compared to a microscope.

#### 3.4.3 Filtration in the cartridge and reading in mCytoCounter

After assessing the membrane and cartridge, the somatic cells were captured in the cartridge and quantified using the mCytoCounter device. The serially diluted somatic cells were externally labeled with CFSE dye and added in the cartridge. Each sample well was loaded with a different dilution of somatic cells. The cartridge was inserted into the device for the device to move the sample through the membrane to the dump. After that, it began reading all the membranes one by one and produced the results in the form of relative fluorescence units. The experiment was performed three times, and the readings were then used to plot a calibration curve, as shown in Figure 6A. The mCytoCounter has a detection range of 10^3^ to 10^6^ SCC/mL, which covers the required cell number range to discriminate between healthy, subclinical, and clinical mastitis.

**Figure 6.**
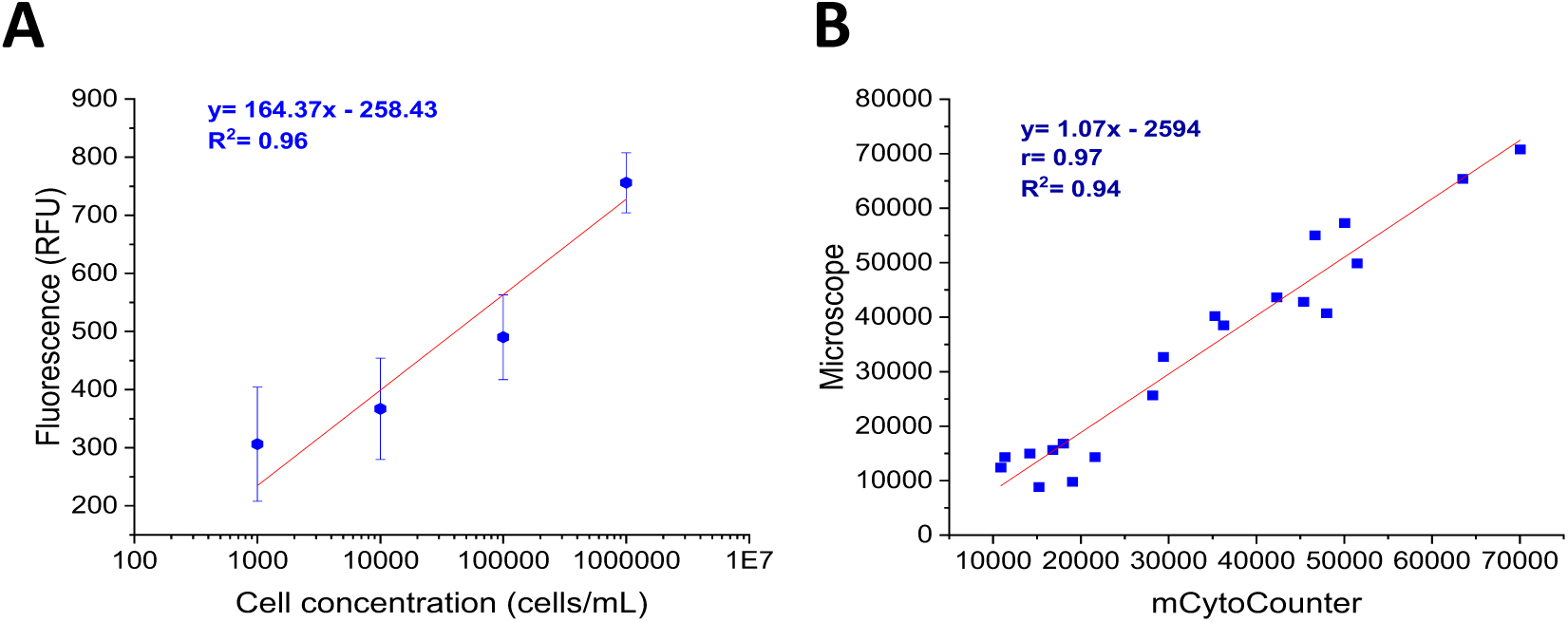
**A-** Calibration plot with mCytoCounter. The experiment was repeated 3 times. **B-** Comparison between mCytoCounter and microscope count. The whole milk was diluted with 1XPBS (1:5).

### 3.5. Benchmarking SCC in mCytoCounter and microscope with farm samples

About 20 fresh milk samples were procured from a dairy farm in New Delhi. The somatic cells were quantified both in the mCytoCounter and the microscope. The milk was diluted (1:5) with 1X PBS and labeled using CFSE dye (90 µM). After labeling, samples were added to the cartridge, and the readings were recorded in the mCytoCounter. The same sample was then added to a flow cell (1cm X 2.8 cm) with a height of 30 µm and visualized in the microscope with a blue filter cube (excitation range 460-495 nm, emission range 510-800nm). The images were captured (20 frames) and cells were counted manually. The RFU value from mCytoCounter and the manual count from the microscope (cells/ frame) were recorded. The calibration plot, generated using the mCytoCounter and the microscope, was used to calculate the exact cell count. The results were plotted and compared in the correlation plot as shown in Figure 6B.

There are many commercially available somatic cell counters such as Lactoscan SCC that use fluorescence microscopy to count the somatic cells (Table 1). The device requires preprocessing, mixing milk with the nucleic acid labeling dye Sofia green. However, its affordability could be a limiting factor in various dairy farms (Chengolova et al.,2021). Similarly, other commercial systems such as DeLaval cell counter and Sceptor2.0, are also very expensive. Sceptor2.0 is a handheld device and requires 6 spin wash cycles of 2 minutes each to remove fats and other interfering substances in the milk. This type of setup is difficult to deploy at the farm level. Nevertheless, these systems offer great reproducibility and can process samples in less than a minute. However, dairy farmers cannot adopt them due to their high cost and bulkiness. On the other hand, our system is an easily deployable, cost-effective system with a high correlation with the direct microscopy count. Therefore, mCytoCounter is a feasible solution to mastitis diagnosis in low-resource settings.

**Table 1.** Comparison between commercial somatic cell counters

| S.No | Name | Principle | Detection Range (cells/mL) | Pre-processing required<sup>a</sup> | Analysis Time (min)<sup>b</sup> | Deployability<sup>c</sup> | Device cost (USD)<sup>d</sup> |
| --- | --- | --- | --- | --- | --- | --- | --- |
| 1. | **Lactoscan SCC(35)** | Fluorescence microscopy | $10^4$ to $10^7$ | Yes, the sample is mixed with | <1 min | ★★ | 2852 |
|  |  |  |  | Sofia green dye |  |  |  |
| 2. | **DeLaval Cell Counter(36 )** | Fluorescence quantification | $10^4$ to $10^7$ | No (Cartridge has Propidium Iodide) | <1 min | ★★★★ | 4086 |
| 3. | **Sceptor 2.0(37)(38)** | Impedance | Up to $1.5 \times 10^6$ | Yes (6 spin wash cycles of 2 min each) | - | ★★ | 7214 |
| 4. | **mCyto-Counter** | Fluorescence quantification | $10^3$ to $10^7$ | Yes, the sample is mixed with CFSE dye | <2 min | ★★★★ | 1000 |

<sup>a</sup> Pre-processing is the steps required to process the samples before it could be added into the device/cartridge
<sup>b</sup> Analysis time considers the time taken by the device to produce results (this doesn't include the pre-processing time)
<sup>c</sup> Deployability parameter considers the weight and size of the equipment. Whether it can be easily moved to the farms and used when required
<sup>d</sup> Device costs of these systems were obtained from various e-commerce websites. Actual cost may vary.

## 4. Conclusion

A quantitative test for diagnosing bovine mastitis has been developed utilizing the fluorescence principle to count somatic cells in milk. The microfluidic cartridge can separate the somatic cells from milk, which are then quantified using a fluorescence detector in the mCytoCounter. The somatic cell quantification results obtained from the mCytoCounter were compared with the count obtained from the fluorescence microscope. The correlation coefficient was 0.97, showing that the mCytoCounter is a rapid and accurate diagnostic method for bovine mastitis. The mCytoCounter has a detection range of 10^3^ SCC/mL to 10^6^ SCC/mL which covers the required cell number range to discriminate between healthy, subclinical mastitis and clinical mastitis. Regardless, future work would evaluate the system with a higher sample size.

Although our present work demonstrates the applicability of mCytoCounter as a diagnostic solution for bovine mastitis, it has the potential to be utilized across a diverse range of samples, such as blood and urine, with slight modifications in the mechanical and electronic designs. The current version of the device demonstrated satisfactory performance under controlled conditions. However, further modifications are underway to enhance its usability and adaptability for on-field applications by farmers.

## CRediT authorship contribution statement

**Neha Khaware:** Methodology, Validation, Formal analysis, Investigation, data curation, Writing-Original draft, Visualization. **Tushar Jeet:** Mechanical design and fabrication. **Prabhu Balasubramanian:** Software, Data curation. **Holger Schulze:** Writing-Review and Editing. **Vivekanandan Perumal:** Writing-Review and Editing, Supervision, Funding acquisition. **Till T. Bachmann:** Writing-Review and Editing, Supervision, Funding acquisition. **Ravikrishnan Elangovan:** Conceptualization, Methodology, Resources, Writing-Review and Editing, Supervision, Project administration, Funding acquisition.

## Acknowledgments

This study is completed as part of the DOSA Project (Diagnostics for One Health and User Driven Solutions for AMR, www.dosa-diagnostics.org). DOSA is jointly funded by the UK Research and Innovation Economic and Social Research Council (grant number: ES/S000208/1), the Newton Fund, and the Government of India’s Department of Biotechnology (grant number: BT/IN/Indo-UK/AMR/03/) and the Ministry of Education (MoE), India under the Grand Challenge Project scheme (MI1798; received by RE and VP) of the Industrial Research & Development Unit, Indian Institute of Technology Delhi. The authors thank Mr. Gaurav Jhakda for procuring milk samples from the dairy farm.

