## Supplemental data for "mCytoCounter, a deployable somatic cell counting device for the assessment of mastitis in dairy farms"

Corresponding author:

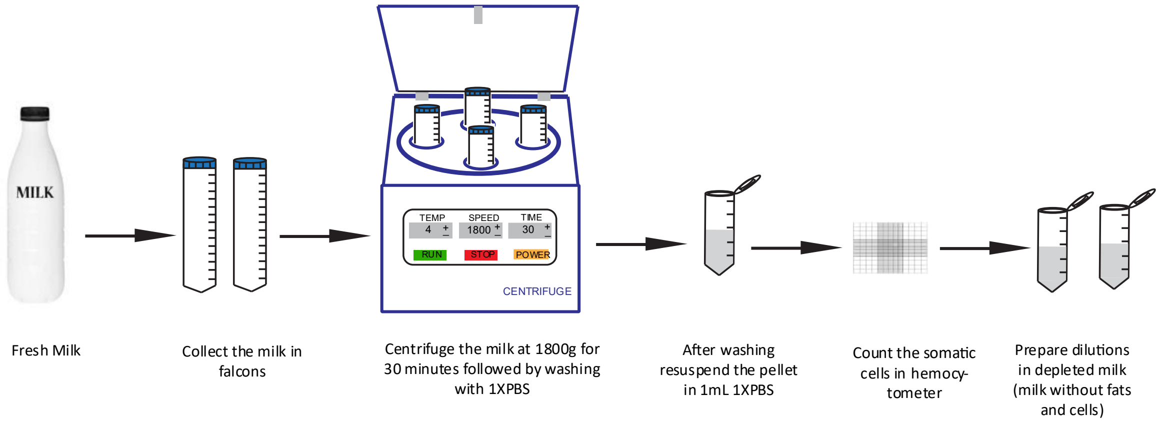


**Figure_S1- Somatic cell isolation protocol.** The fresh milk is centrifuged and washed, and the final pellet is resuspended in 1XPBS. The isolated somatic cells are then counted using a hemocytometer, and dilutions are made in the depleted milk (milk from which the cells have been removed).

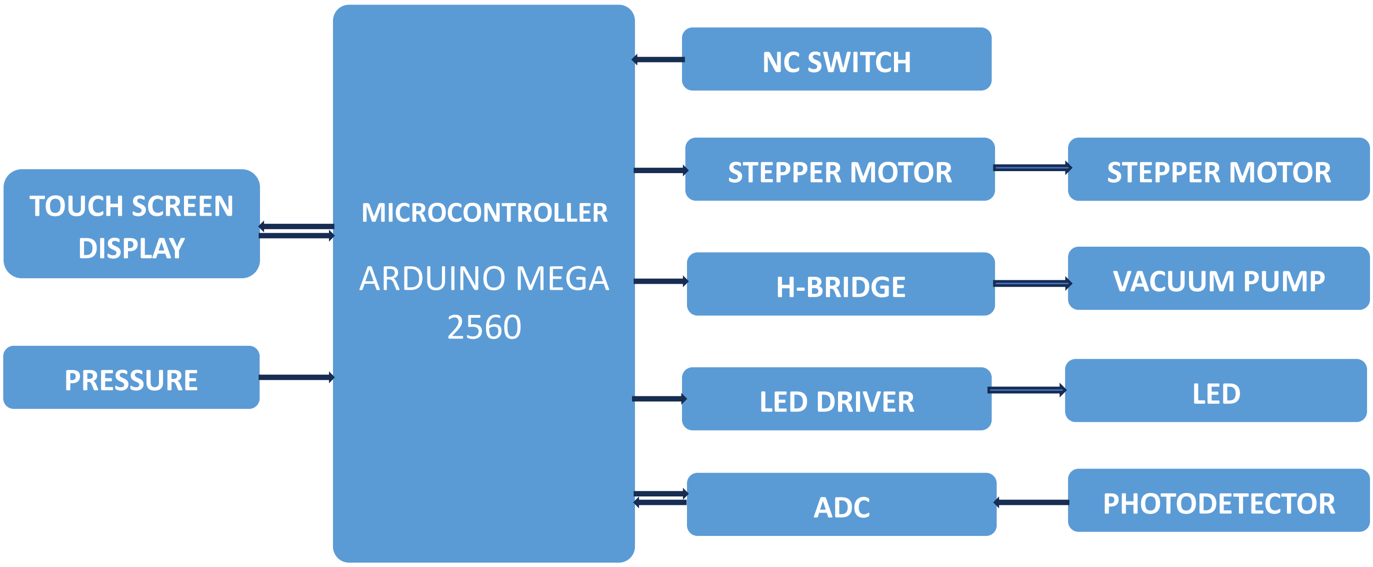


**Figure_ S2- Block diagram of Printed circuit board.** The microcontroller, an integrated circuit, interacts with the Graphical User Interface via touchscreen pad and initiates the program. It controls the motor to move the photodetector, which collects the emission signals from the sample; the vacuum pump, which controls the liquid flow; the LED, which excites the sample; and the photodetector, which collects the fluorescence signals. The signals are converted into relative fluorescence units and displayed on the screen.

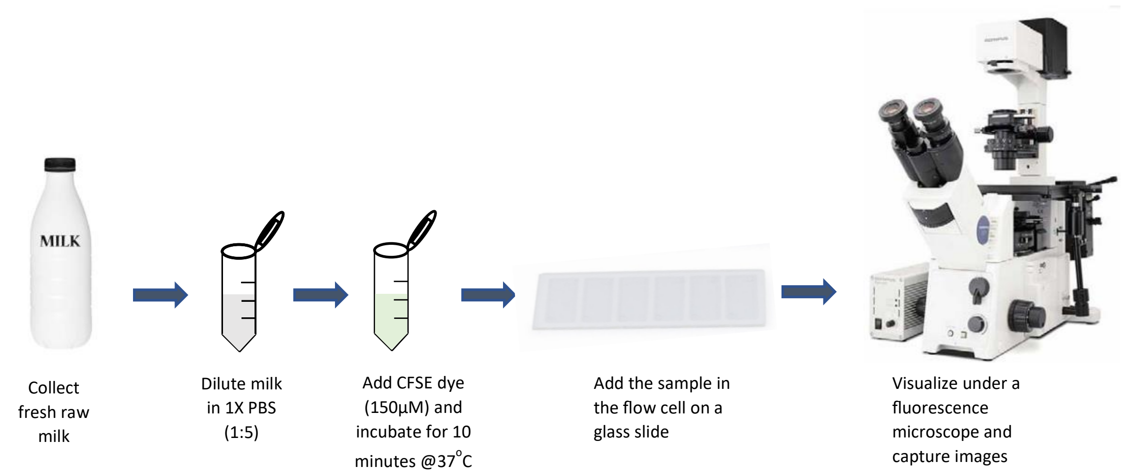


**Figure_S3- Protocol for direct milk microscopy.** The fresh milk is diluted with 1X PBS and somatic cells are labelled with CFSE dye. The sample is then added to the glass slide with flow cell to visualize under the fluorescence microscope.

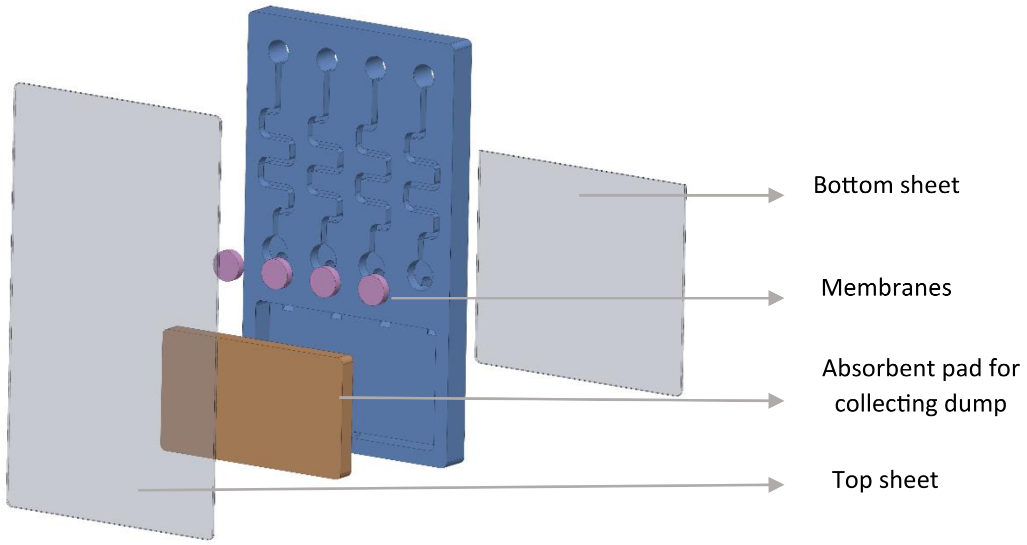


**Figure_S4- Microfluidic cartridge assembly.** The 3D-printed cartridge is assembled with GF/B membranes and an absorbent pad to collect the dump. Finally, the cartridge is sealed with a transparent sheet on the front and back.

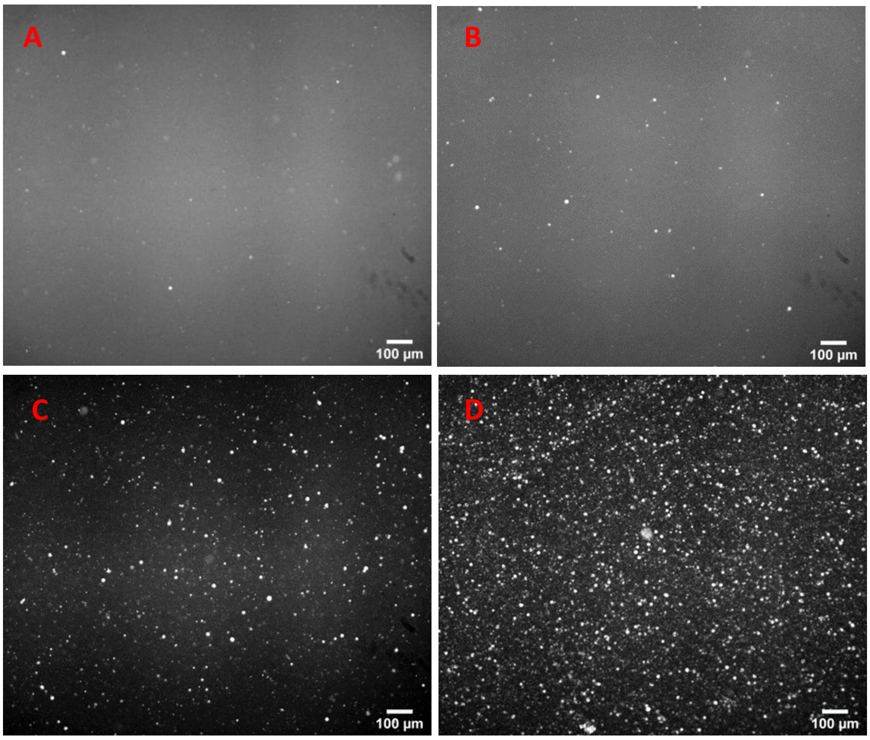


**Figure_S5- Somatic cells visualization under inverted fluorescence microscope (Olympus IX71).** Somatic cells were serially diluted (with 1XPBS) and labeled with CFSE dye (90µM). A= $10^4$ SCC/mL, B=$10^5$ SCC/mL , C= $10^6$ SCC/mL and D= $10^7$ SCC/mL. The somatic cells were visualized in a flow cell (1cm X 2.8cm X 0.003cm)
